# Can endogenous fluctuations persist in high-diversity ecosystems?

**DOI:** 10.1101/730820

**Authors:** Felix Roy, Matthieu Barbier, Giulio Biroli, Guy Bunin

## Abstract

When can complex ecological interactions drive an entire ecosystem into a persistent non-equilibrium state, where species abundances keep fluctuating without going to extinction? We show that high-diversity spatially-extended systems, in which conditions vary somewhat between spatial locations, can exhibit chaotic dynamics which persist for extremely long times. We develop a theoretical framework, based on dynamical mean-field theory, to quantify the conditions under which these fluctuating states exist, and predict their properties. We uncover parallels with the persistence of externally-perturbed ecosystems, such as the role of perturbation strength, synchrony and correlation time. But uniquely to endogenous fluctuations, these properties arise from the species dynamics themselves, creating feedback loops between perturbation and response. A key result is that the fluctuation amplitude and species diversity are tightly linked, in particular fluctuations enable dramatically more species to coexist than at equilibrium in the very same system. Our findings highlight crucial differences between well-mixed and spatially-extended systems, with implications for experiments and their ability to reproduce natural dynamics. They shed light on the maintenance of biodiversity, and the strength and synchrony of fluctuations observed in natural systems.

---

While large temporal variations are widespread in natural populations [1, 2], it is difficult to ascertain how much they are caused by external perturbations, or by the ecosystem’s internal dynamics, see e.g. [3, 4]. In particular, both theoretical tools and empirical results come short of addressing a fundamental question: can we identify when fluctuations in species abundances arise from complex ecological interactions?

Our focus here is on high-diversity communities. Historically, studies of endogenous fluctuations have focused on single populations or few species [5, 6]. On the other hand, theories of many-species interaction networks often center on ecosystems that return to equilibrium in the absence of perturbations [7]. Some authors have even proposed that endogenous fluctuations are generally too rare or short-lived to matter, since they can be self-defeating: dynamics that lead to large erratic variations cause extinctions, leaving only species whose interactions are less destabilizing, with extinctions continuing until an equilibrium is reached [8, 9].

Many-species endogenous fluctuations can only persist if they do not induce too many extinctions. Extinction rates are related to the amplitude of fluctuations [10, 11], their synchrony [12] and their correlation time [13]. The peculiarity of endogenous fluctuations is that these properties arise from the species dynamics, and therefore feed back on themselves. A theory of these feedbacks is however lacking.

Here we propose a novel quantitative approach, and show that many-species endogenous fluctuations can persist for extremely long times. Furthermore, they can be realized in experimental conditions, and identified in these experiments by multiple characteristic features. Crucially, we show that states with higher species diversity have stronger fluctuations, and vice versa. We also offer reasons why they may not have been observed in previous studies, and directions in which to search. An important factor in maintaining a dynamically fluctuating state is the spatial extension of the ecosystem, here modeled as a metacommunity: multiple patches (locations in space) that are coupled by migration.

Our strategy is the following. We first propose and simulate experiments to show that persistent fluctuations can be very elusive in a single well-mixed community, yet attainable in a metacommunity via three main ingredients: the existence of multiple patches, moderate migration fluxes coupling them, and differences in conditions between patches. These three ingredients can mitigate the likelihood that large fluctuations within a patch will lead to overall extinctions (see Fig. 1), and make it possible for species to persist in highly fluctuating states.

**Figure 1.**
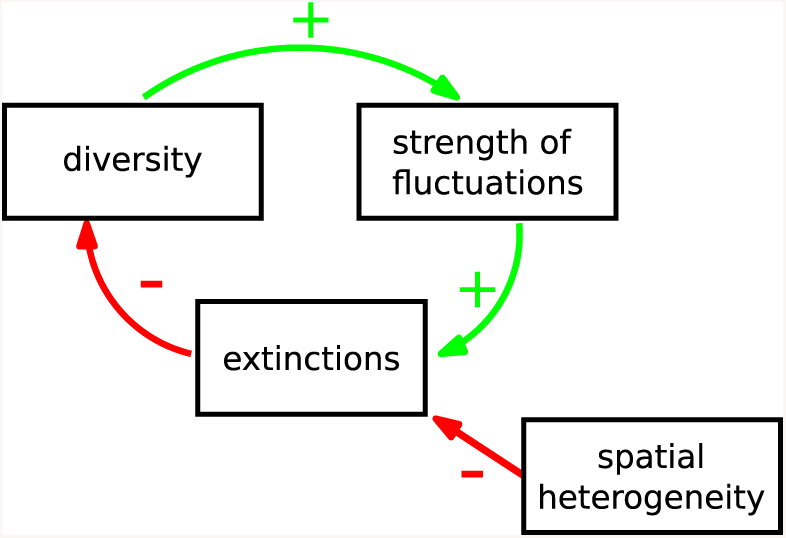
The fluctuation-diversity feedback cycle. Species diversity is required to maintain endogenous fluctuations. But these fluctuations cause extinctions, which reduce diversity. This negative feedback cycle can lead to the disappearance of endogenous fluctuations, especially in a well-mixed community. However, if spatial heterogeneity can limit extinctions, this negative feedback loop may slow down and create a fluctuating state that persists for very long times.

We then offer a quantitative understanding of this phenomenon. We build on the analytical framework developed in [14] (dynamical mean-field theory) that allows us to investigate, in a quantitative and predictive way, the conditions under which robust fluctuations can arise from complex interactions. This theory exactly maps a deterministic metacommunity (many-species dynamics over multiple spatial locations) to a *stochastic representative metapopulation* (single-species dynamics over multiple spatial locations). It predicts the distribution of abundance, survival and variability for a species subjected to “noise” that results from other species in the same community, rather than external perturbations. Dynamical mean-field theory allows us to analyze these fluctuations, and show that the effective stochasticity of species dynamics is a manifestation of high-dimensional chaos.

The intuitive picture that emerges from our analysis is the following: the persistence of endogenous fluctuations, which can be found in a wide range of realistic conditions, requires a balancing act between forces that stabilize and destabilize the dynamics, see Fig. 1. On the one hand, the system needs to preserve a high diversity (both in terms of species number and interaction heterogeneity), as it is known [7, 15] that lower diversity leads to a stable equilibrium. On the other hand, the system also has to limit excursions towards very low abundances. This requires weeding out species that induce unsustainable fluctuations, and rescuing the others from sudden drops. To accomplish that, the system relies on asynchronous dynamics between different spatial locations, and finite strength and correlation time of the abundance fluctuations.In addition, despite large fluctuations in the abundances of all species, that strongly affect the species growth in any given patch, long-lasting “sources” emerge for some of the species, i.e. patches where these species are more likely to remain away from extinction. Rare dynamical fluctuations leading to extinction in a given patch are hampered by migration from the other patches, which keeps the system in a non-equilibrium state. We show that, with moderate migration and some spatial heterogeneity, high-diversity dynamical states can be reached where species populations fluctuate over orders of magnitude, yet remain bounded for very long times above their extinction threshold.

Our findings allow us to paint a more precise picture of when persistent endogenous fluctuations can arise. We conclude with a discussion of the implications for biodiversity and ecosystem stability, and predictions for future experiments on community dynamics.

## Proposed experiments

In the following, we introduce our results via a set of proposed experiments, realized in simulations, see Fig. 2. These results are later explained in the theoretical analysis. All parameters for simulations are detailed in Appendix A.

**Figure 2.**
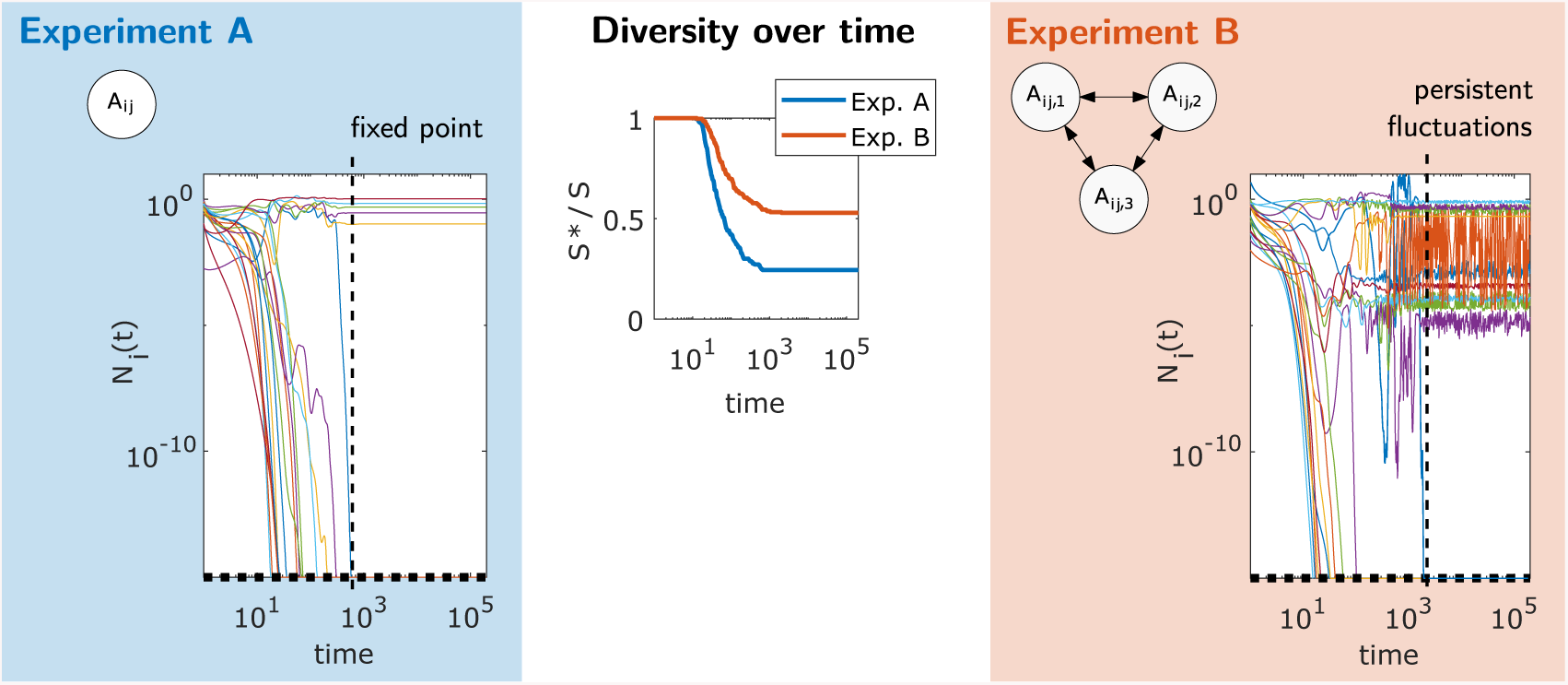
Numerical realization of the proposed experiments, illustrating conditions that lead to a fixed point or persistent fluctuations. (A) A single patch (well-mixed community) with an interaction matrix *A*_*ij*_. (B) Multiple patches connected by migration, with slightly different conditions (e.g. temperature or resources) in each patch, represented here by location-dependent parameters such as *A*_*ij,u*_. In the right and left panels we show the time evolution of a few representative species abundances *N*_*i*_(*t*): Experiment A, with a single patch (*M* = 1) reaches a fixed point, while in experiment B a meta-community with *M* = 8 patches reaches a stationary chaotic state (*S* = 250). Middle panel: Fraction of persistent species (*S*\* out of a pool of *S* = 250 species) as a function of time.

We focus on a meta-community which consists of *M* patches (well-mixed systems) connected by migration, and isolated from the external world. We consider generalized Lotka-Volterra equations for the dynamics of the abundance *N*_*i,u*_ of species *i* in patch *u*:

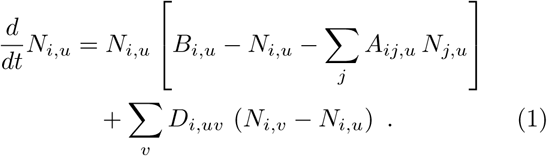

where *A*_*ij,u*_ are the interactions coupling the species, *B*_*i,u*_ represents the equilibrium abundance in absence of interactions and migration (known as the carrying capacity), and *D*_*i,uv*_ are the migration rates between patches *u* and *v* In addition, an extinction threshold is implemented as follows: when a species’ abundance goes below a cut-off *N*_*c*_ in *all* patches, the species is removed from the metacommunity and cannot return^1^. This threshold corresponds to the minimum sustainable number of individuals, hence 1*/N*_*c*_ sets the scale for the absolute population size (*P*) of the species. For simplicity, we take *D*_*i,uv*_ = *d/* (*M* − 1) and *N*_*c*_ identical for all species and patches.

The species are assumed to have unstructured interactions (e.g. they belong to the same trophic level), meaning that *A*_*ij,u*_ are sampled independently^2^ and identically for different (*i, j*). For a given species pair, its interactions *A*_*ij,u*_ vary somewhat with *u*; this variability corresponds to small differences in the conditions between the patches [16]. In the simulation examples we set all carrying capacities *B*_*i,u*_ = 1; the phenomena described below are also found if *B*_*i,u*_ vary between patches in addition to, or instead of the interaction coefficients.

Our proposed experiments, illustrated by dynamical simulations, are the following:

A. First, we model a single patch, *M* = 1 initially containing *S* = 250 species^3^. Species go extinct until the system relaxes to a fixed point (stable equilibrium), see left panel of Fig. 2.
B. We now take *M* = 8 patches with the same initial diversity *S* = 250 and interaction statistics as in (A). For each pair of interacting species, *A*_*ij,u*_ varies slightly with location *u*, with a correlation coefficient *ρ* = 0.95 between patches. The abundances now fluctuate without reaching a fixed point, see right panel of Fig. 2. At first the diversity decreases as species go extinct, but this process dramatically slows down, and the diversity is unchanged at times on the order of 10^5^, see middle panel of Fig. 2.

Three essential observations emerge from simulating these experiments, and repeating them for different parameters. First, species diversity and the strength of endogenous fluctuations are tightly bound, each contributing to the other’s maintenance. Second, as shown in Fig. 2, species trajectories first go through a transient phase where they fluctuate over many orders of magnitude, causing numerous extinctions which lead to a reduction of variability, until a fixed point (for *M* = 1) or non-equilibrium state (for *M* = 8) with weaker fluctuations is reached. Third, the qualitative difference between experiments A and B is robust to changes in parameter values. Changes in *N*_*c*_ and *d* affect only quantitatively the states that are reached in experiment B, see Fig. 3(top). For instance, by increasing the population size *P* = 1*/N*_*c*_, we can reach dynamically fluctuating states with higher long-time diversities, as shown in Fig. 3(bottom). When the population size is reduced by increasing *N*_*c*_, the long-time diversity decreases, but remains high until *N*_*c*_ ∼ 10^−2^ 10^−1^, where it decreases dramatically. For example, the diversity shown in Fig. 2(right) is 80% _*±*13%_ higher than that reached for fixed-points at higher *N*_*c*_. Similarly, as long as the migration coefficient is in the range *d* ≲ 0.1 the main qualitative results remain unaltered.

**Figure 3.**
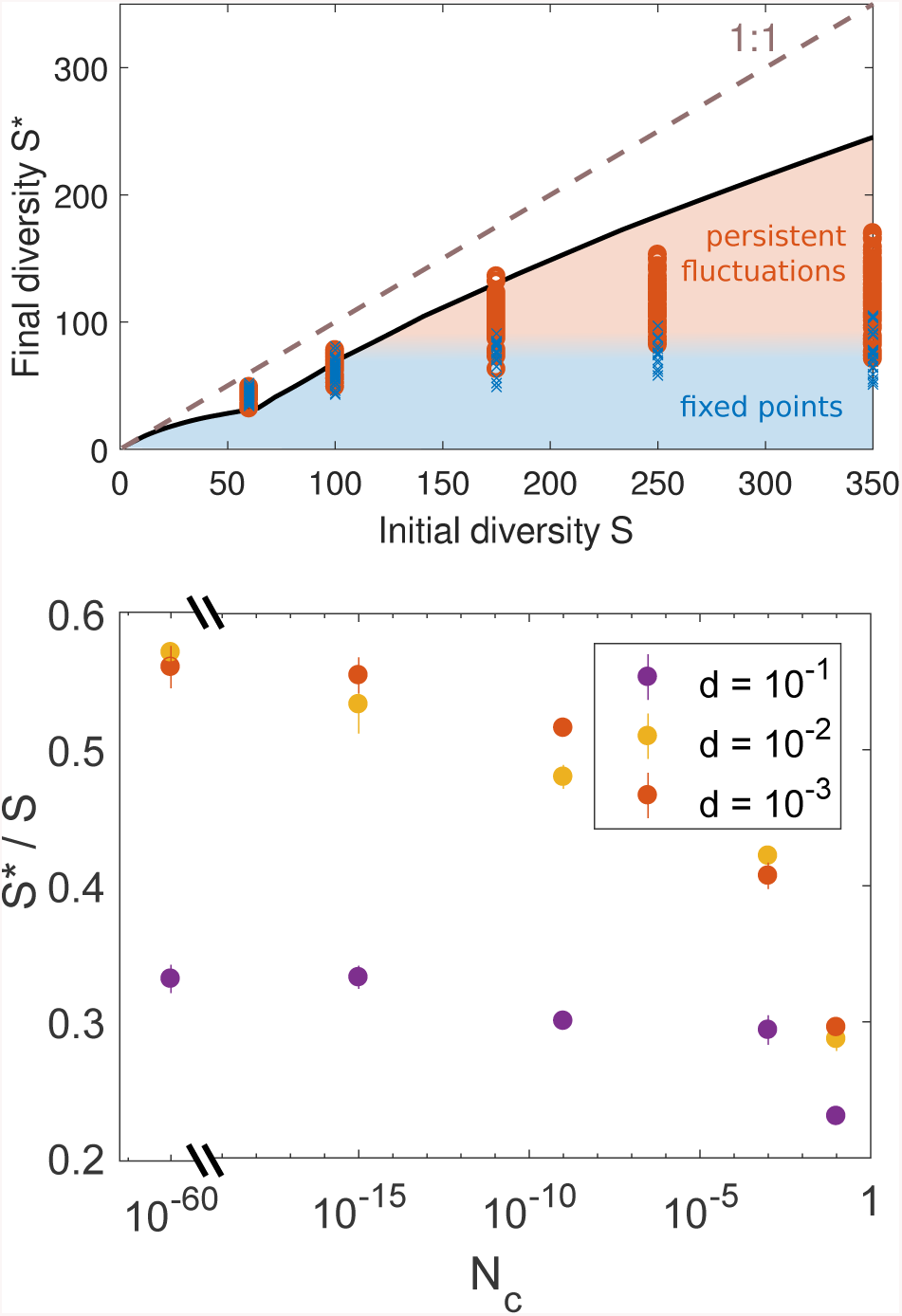
Top: Species diversity at long times, compared to the theoretical bound obtained in Appendix D for large *S* (solid line). The bound depends on the distribution of interactions, carrying capacities and initial pool size *S*. Each symbol represents the state at the end of one simulation run, with fluctuating states (circles) and fixed points (crosses). States closer to the theoretical bound (with higher diversity) also exhibit larger fluctuations and are more difficult to reach due to extinctions in the transient dynamics (see Fig.2). The dashed line represents full survival (*S*\* = *S*). Bottom: The final diversity is set by the transient dynamics, which is affected by factors such as the migration strength and the total population size (1*/N*_*c*_).

## Theory

We now aim to understand which conditions allow a fluctuating state to be reached and maintained without loss of species.

### Dynamical Mean Field Theory

We build on a powerful theory, known as Dynamical Mean Field Theory (DMFT) that exactly maps the deterministic meta-community problem (many species in multiple patches) to a stochastic meta-population problem (single species in multiple patches). When species traits and interactions are disordered, e.g. drawn at random from some probability distributions, all species can be treated as statistically equivalent [17]. We can then describe the whole system by following the trajectory of a single species, randomly sampled from the community, and studying its statistics. In the DMFT framework, the effect of all other species on that single species is encapsulated by an “ecological noise” term generated by their fluctuations. This is analogous to the use, in physics, of thermal noise to represent interactions between an open system and its environment. Since species are statistically equivalent, the properties of this ecological noise can be self-consistently obtained from the dynamics of the single species.

While the theory applies to all times [14], as discussed in Appendix B, we only consider here the stationary state^4^ reached after a long time, in which extinctions are already rare. In that state, observables such as the mean abundance are stable over time, and two-time measures, such as correlation functions between times *t* and *t*′, depend only on the difference *t* − *t*′. This entails that each species abundance fluctuates with a finite correlation time, i.e. it tends to return to some constant characteristic value after a finite time.

The result of this mapping is that the abundance *N*_*u*_ of a given species in patch *u* undergoes stochastic dynamics,

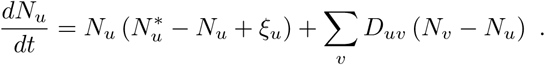

This equation models the dynamics of the target species, including its interactions with other species whose abundances are fluctuating. The contribution of interactions can be separated into a time-independent and a time-dependent parts. The time-independent part goes into 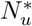, the characteristic value around which the species abundance will fluctuate in the patch. It differs between species and between patches, due to interactions and to environmental preferences modeled by *B*_*i,u*_ in Eq. (1), and follows a multivariate Gaussian distribution. The time-dependent part is encapsulated in *ξ*_*u*_ (*t*), a Gaussian noise with a finite correlation time.

As the quantities 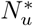 and *ξ*_*u*_ (*t*) result from interaction with other species, which are statistically equivalent to the target species, one can express their properties from the statistics of *N*_*u*_(*t*) itself, as shown in Appendix B. The most important features are that time-dependent Gaussian noise *ξ*_*u*_ (*t*) has zero mean and a co-variance *C*_*ξ*_ (*t, t*′), which is directly related^5^ to the time auto-correlation *C*_*N*_ (*t, t*′) of *N*_*u*_ (*t*) within a patch, and to *σ*^2^ = *cS* var (*A*_*ij*_) the variance of interactions rescaled as in [7]. Moreover, the covariance of the 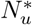 is fixed by the time auto-correlation of *N*_*u*_ (*t*), both within and in-between patches. In principle, the noise is also correlated between patches, but this is a small effect in the dynamical regime of interest to us, see the next section. Note that *C*_*ξ*_ (*t, t*′) vanishes when a stable equilibrium is reached.

The analysis of the DMFT equations clarifies the main effect of coupling patches by migration: patches with higher 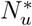 tend to act as sources, i.e. the species most often grows there, and migrates out to sites where it cannot grow (sinks). We show directly from simulations of the Lotka-Volterra equations in Fig. 4 that species have particular patches which tend to act as sources consistently over long times. This fact is counter-intuitive, as the abundances of all species may be fluctuating over orders of magnitude in any given patch, yet this patch will retain its identity as a source (or sink) when averaging over long time periods. The variability of the 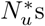 between patches thus leads to an insurance effect, since it is enough to have one patch acting as a source to avoid extinction of the species in the others.

**Figure 4.**
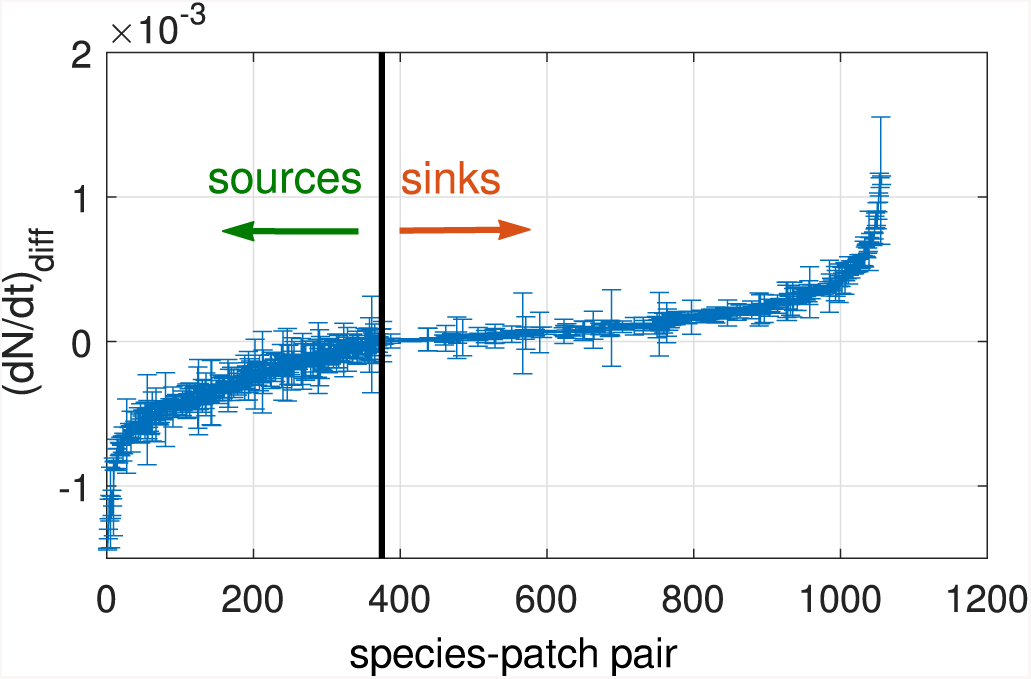
Sources maintain their identity over time. The degree to which a patch is a source for a given species is measured by (*dN/dt*)_diff_, the contribution of diffusion to the change of *N* (*t*), which is negative for sources and positive for sinks. We show all species-patch pairs ordered by the average of this quantity over long times, with error bars giving its standard deviation. For 94% of sources, and 85% of all species-patch pairs, this quantity (*dN/dt*)_diff_ retains its sign most of the time, being at least one standard deviation away from zero.

### Reaching and maintaining a dynamical state

Let us first consider a single community (*M* = 1). For a species to survive for long periods of time, it follows from DMFT that it must have positive *N*\*, or else *N* (*t*) decays exponentially until the species goes extinct. Even if *N*\* > 0, there is still a probability (per unit time) of extinction, which depends on *N*\*, *N*_*c*_ and on the strength of the noise *ξ* (*t*). Following extinctions, a remaining species interacts with fewer fluctuating other species, causing the strength of the noise to decrease, and with it the probability for extinction, see feedback loop in Fig. 5.

**Figure 5.**
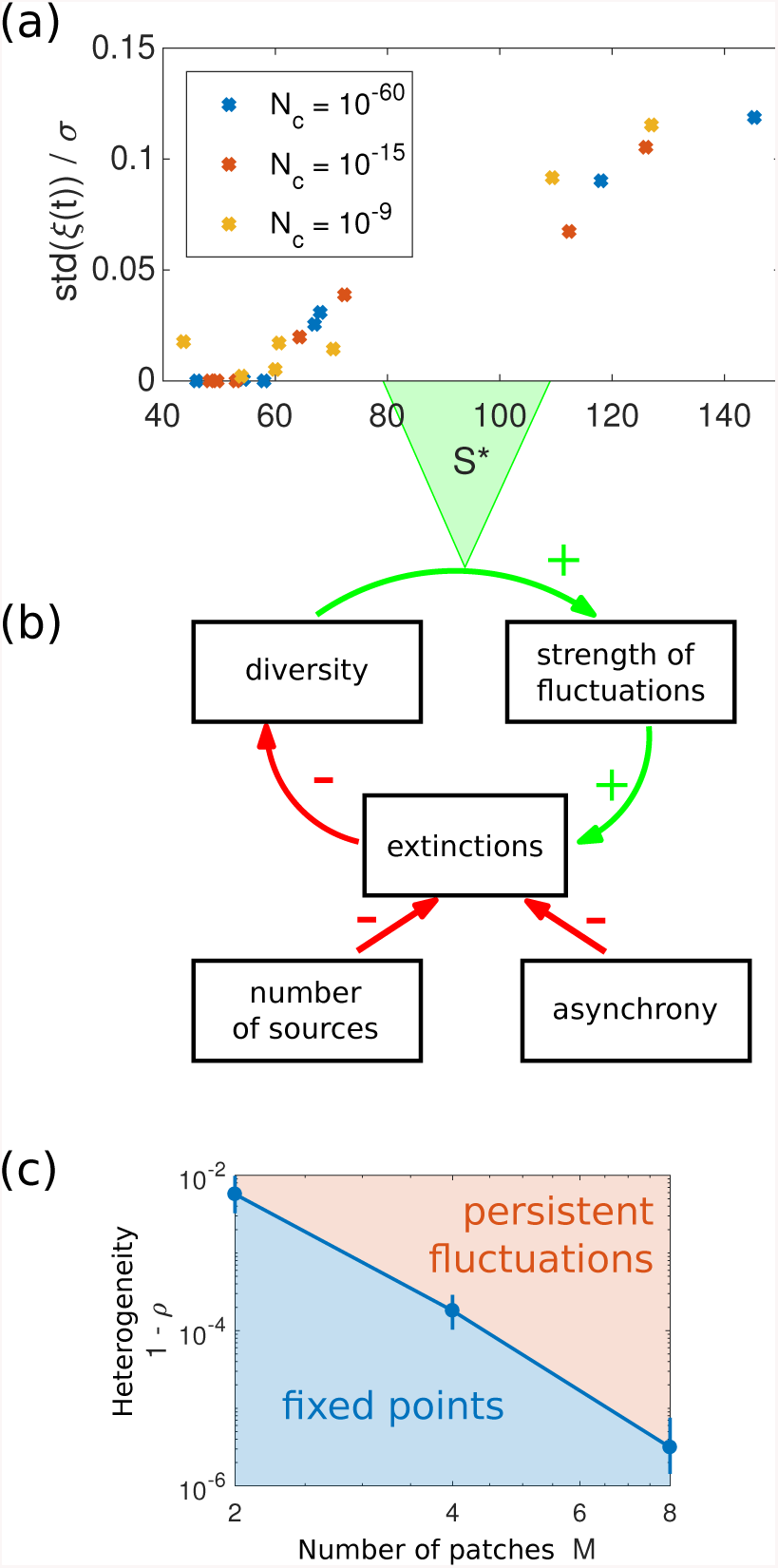
Revisiting the noise-diversity feedback cycle in the light of our theoretical framework. (a) Quantitative relationship between species diversity *S*\*, i.e. the number of coexisting species, and strength of fluctuations std(*ξ*) for *M* = 8 (rescaled by interaction heterogeneity *σ*). (c) Patch number *M* and heterogeneity 1 − *ρ* (defined from the correlation coefficient *ρ* between interactions *A*_*ij,u*_ in different patches *u*) both contribute to the persistence of endogenous fluctuations by two means, shown in (b): they create source patches where a given species will tend to grow (see Fig.4), and allow the asynchrony of fluctuations in different patches. These two factors mitigate the likelihood that endogenous fluctuations will induce species extinctions and cause their own suppression.

We can develop an analytical treatment for very small cut-off *N*_*c*_ (large population size). In this case there is a large difference in time-scales between the short-term dynamics induced by endogenous fluctuations, and the long-term noise-diversity feedback cycle discussed above. In fact, the extinctions driving this feedback are due to rare events in which the abundance of species with a positive *N*\* decreases below the (very small) cut-off *N*_*c*_. For a species in an isolated patch (*M* = 1), the time-scale for such an event is known [10, 11] to be of order of *τ* (1*/N*_*c*_)^*a*^ where *τ* is a characteristic time of the endogenous fluctuations, and *a* = 2*N*\**/W* is independent^6^ of *N*_*c*_, with *W* the amplitude of the endogenous fluctuations,

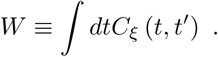

The important point here is that, although endogenous fluctuations disappear eventually, there is a clear *separation of time-scales* between typical endogenous fluctuations, that are fast and lead to a quasi-stationary dynamical state, and rare extreme fluctuations that cause extinctions and push the ecosystem into a different state.

While a single community might in principle achieve long-lasting endogenous fluctuations, this however requires unrealistically large population sizes and species number, see Appendix E. Migration between multiple patches substantially enhances persistence due to the spatial insurance effect [12]: species are more unlikely to go extinct because they need to disappear everywhere at once. The time scale for such an event is 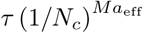 with 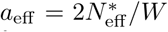 where 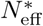 is an effective value for *N*\* of a species across patches, an expression for which is given in Appendix C. This result is thus similar to the one identified above for one patch, raised to the power *M*. These results assume that *W* is finite, and that the noise acting on a species is independent between patches (asynchrony). Indeed, for moderate values of *D* and *ρ* not too close to one, simulations show that *C*_*ξ*_ (*t, t*′) is a well-behaved function of *t* so that *W* is finite, and the correlation between patches is found to be very small, see Appendices B,C.

These expressions provide a quantitative description of the feedback cycle in Fig. 5. Endogenous fluctuations disappear on the time scale at which species with characteristic abundance 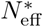 of order one would go extinct. We must further account for the vanishing strength of the noise *W* as species disappear. This is shown in Fig. 5(a), where the strength of the fluctuations is tightly linked to species diversity, and is zero at the diversity of fixed points. Hence, extinctions significantly increase *a*_eff_, reducing the chance for further extinctions.

This picture agrees well with the analytical predictions, which can be obtained for very small but positive *N*_*c*_ and *D* by DMFT. These give the distribution of 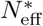, whose integral over positive 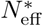 amounts to the maximal total diversity at long times. We show in Fig. 6 that extinct species are generally those that have lower values of 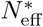: we compare the analytically predicted distribution of 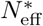 to the one observed in simulations, and see that most missing species had low values of 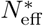. The difference becomes smaller for lower *N*_*c*_. Due to these differences, the obtained diversities are lower than the theoretical maximum, see Fig. 3(top).

**Figure 6.**
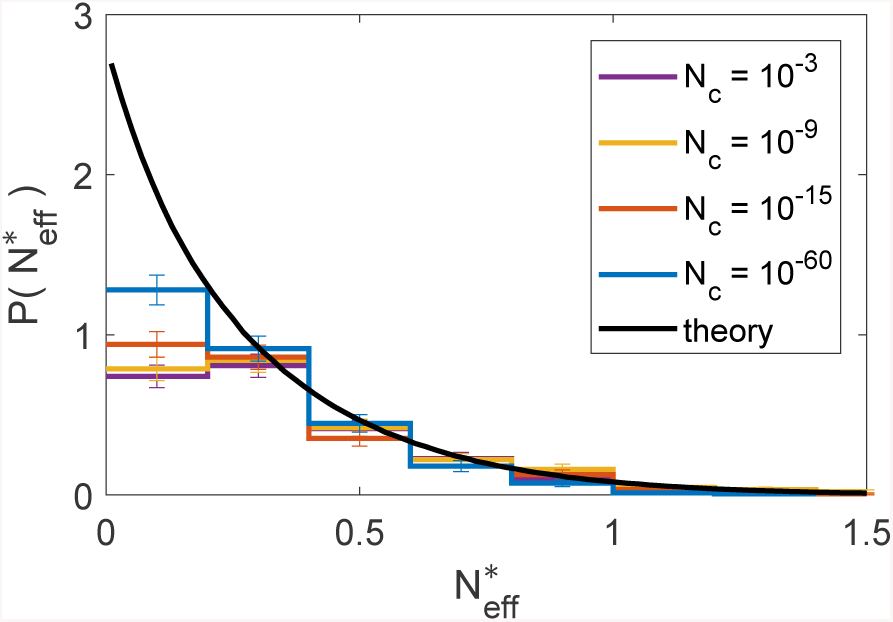
Distributions of the characteristic abundance 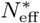 of surviving species, compared with the theoretical prediction for maximal diversity, showing that the lower diversity in simulations is mostly due to losing species with lowest 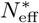 (left-most bin). Reducing *N*_*c*_ (increasing population size) affects diversity mainly by allowing these “rare” species to persist.

As stressed above, the asynchrony of fluctuations in different patches is crucial: it allows some species to survive with positive characteristic abundance 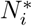 in at least one of the patches. This leads to a higher total number of long-term persisting species, decreases the likelihood of fluctuations to small abundances, and hence increases the stability of a dynamically fluctuating state.

If the migration rate *D* is too strong or *ρ* very close to one, dynamics in the different patches synchronize, quickly annulling the insurance effect. However, minor (few percent) changes in interaction coefficients or carrying capacities between patches are enough to maintain this effect, see Fig. 5(bottom); we don’t need to impose coexistence artificially, e.g. by requiring that every species has at least one refuge (a patch so favorable to it that it always dominates there). These little variations in the interaction coefficients are highly plausible, as interaction strength can vary with many factors, including resource availability [18], or temperature and its influence on metabolism [19]. The heterogeneity *ρ* required to reach a fluctuating state decreases with *M*, see Fig. 5(bottom).

In practice, maintaining a dynamical state seems unfeasible for only one patch, at least for reasonable values of population size *P* = 1*/N*_*c*_ and species number *S*, see discussion in Appendix E. Yet the combined effect of the two phenomena described above allows for very long-lived endogenous fluctuations in metacommunities, already for *M* = 2 patches.

## Conclusions

Species interactions can give rise to long-lasting fluctuating states, which both require and allow the maintenance of high species diversity. This can happen under a wide range of conditions, which we have illustrated in simulated experiments, and identified through an analytical treatment based on Dynamical Mean-Field Theory. While we have drawn parallels with the theory of stability and coexistence in externally-perturbed ecosystems [6, 10–12], our approach also highlights essential differences between environmentally-driven and endogenous fluctuations. We show that many-species dynamics induce feedback loops between perturbation and response, and in particular a tight relationship between fluctuation strength and species diversity, which are absent from externally-perturbed ecosystems. Moreover, while similar species can display correlated responses to environmental stochasticity [20], we expect here that their trajectories will be starkly different and unpredictable, due to high-dimensional interactions which lead to complex dynamics. The resulting picture from DMFT is that the abundance of any given species undergoes stochastic dynamics with a finite correlation time. This means that the trajectory of the species abundance cannot be predicted after a time that is large compared to the correlation time–a hallmark of chaos, also found in other models of high-dimensional systems [21]. Our theory paves the way for quantitative testing of these fingerprints of diversity-driven fluctuations in data.

In a counterpoint to classic results [7], we have shown that, while highly diverse ecosystems are unstable, they might still persist: extinctions can be avoided and biodiversity maintained, despite species abundances fluctuating over multiple orders of magnitude. We do observe a negative feedback loop, in which endogenous fluctuations cause extinctions, and eventually lead to their own disappearance as the ecosystem reaches a lower-diversity stable equilibrium. But this self-suppression of fluctuations can be mitigated by a number of factors, among which space is particularly important.

In a single well-mixed community, we expect that persistent fluctuations might not be observed in practice: while theoretically possible, they may require unrealistic population sizes and species numbers. But spatial extension and heterogeneity can dramatically reduce these requirements, in a way that parallels the insurance effect against exogenous perturbations. When fluctuations are not synchronized across space, some patches can act as sources, from which failing populations will be rescued through migration [6, 12]. Here, we find that the existence of sources is surprisingly robust: even if there is no location where the environment is favorable to a given species, source patches can arise from interactions, and endure for long times despite the large fluctuations in species abundances. By allowing fluctuations without extinctions, spatial heterogeneity helps maintain species diversity, and thus the fluctuations themselves. This result is robust over a wide range of parameters, as it only calls for moderate values of inter-patch migration: the rate *D* must be such that, over the typical time scale of abundance fluctuations, many individuals can migrate out of a patch (allowing recolonization in the absence of global extinction), while representing only a small fraction of the population in that patch.

A crucial result is that this condition suffices to ensure that synchronization between patches is absent, and that the total strength and correlation time of the noise within patches (*W* above) remain bounded for finite populations and finite migration rates between patches. This is in contrast to alternative scenarios where noise correlations decay slowly with time [22]. This result is non-trivial for endogenous fluctuations, as the existence of feedbacks (encoded in the self-consistent equations of the DMFT framework) can potentially lead to synchronization and long-time correlations in the noise. Yet we demonstrate that synchrony is avoided, both through direct simulations, and by building an analytical theory based on these assumptions, whose predictions match simulations quantitatively.

In conclusion, non-equilibrium fluctuating states might be much more common than suggested by experiments and theory for well-mixed communities. And since these fluctuations permit the persistence of more species than could coexist at equilibrium, we might also expect significantly higher biodiversity in natural environments.

## Acknowledgments

It is a pleasure to thank J.-F. Arnoldi, J.-P. Bouchaud, C. Cammarota, D. S. Fisher and M. Loreau for helpful discussions. G. Bunin acknowledges support by the Israel Science Foundation (ISF) Grant no. 773/18. M. Barbier was supported by the TULIP Laboratory of Excellence (ANR-10-LABX- and by the BIOSTASES Advanced Grant, funded by the European Research Council under the European Union’s Horizon 2020 research and innovation programme (666971). F. Roy acknowledges support by Capital Fund Management - Fondation pour la Recherche. G. Biroli was partially supported by the Simons Foundation collaboration Cracking the Glass Problem (No. 454935).

## Appendix A: Model parameters used in simulations

For convenient reference, this Appendix includes the parameters for all simulations. The model is given in Eq. (1). All *B*_*i,u*_ = 1 and all *D*_*i,uv*_ = *d/* (*M* − 1). The *A*_*ij,u*_ are independent for different (*i, j*) pairs (except in Appendix 13).

In Fig. 2, the probability of *A*_*ij,u*_ to be non-zero is *c* = 1*/*8, and the non-zero elements are sampled from a normal distribution with mean (*A*_*ij,u*_) = 0.3, std (*A*_*ij,u*_) = 0.45. The same elements *A*_*ij,u*_ are non-zero across all patches *u*. The correlation coefficient between non-zero *A*_*ij,u*_ in different patches is *ρ* = corr [*A*_*ij,u*_, *A*_*ij,v*_] = 0.95 for *u* ≠ *v*. (The correlation is 0.964 when interactions with *A*_*ij,u*_ = 0 are also counted.) The initial (pool) diversity is *S* = 250. In Fig. 2(A), *M* = 1. In Fig. 2(B), *M* = 8 patches and *d* = 10^−3^. The cutoff is *N*_*c*_ = 10^−15^. Fig. 3(bottom), uses the same parameters as Fig. 2, but with a range of values for *d, S* and *N*_*c*_.

Fig. 4 uses the runs shown in Fig. 2(B). Standard deviation and mean are estimated from 1601 time points during the time period *t* = [10^4^, 2 · 10^5^]. Fig. 6 uses multiple runs, with the same parameters as 2, except for *d* = 10^−4^ and the values of *N*_*c*_ that are detailed in the figure legend.

Fig.5(top) uses the same parameters as 6. Fig. 5(bottom), shows the line where half of the runs are fixed points, and half continue to fluctuate until *t* = 2 · 10^5^.It uses same parameters as 6, except with *D* = *d/* (*M* − 1) = 10^−4^.

In Fig. 5(top), the size of the fluctuations are calculated from 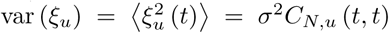, with 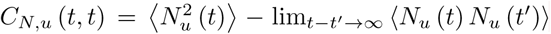. For more details on the averaging, see Appendix B, Fig. 7.

**Figure 7.**
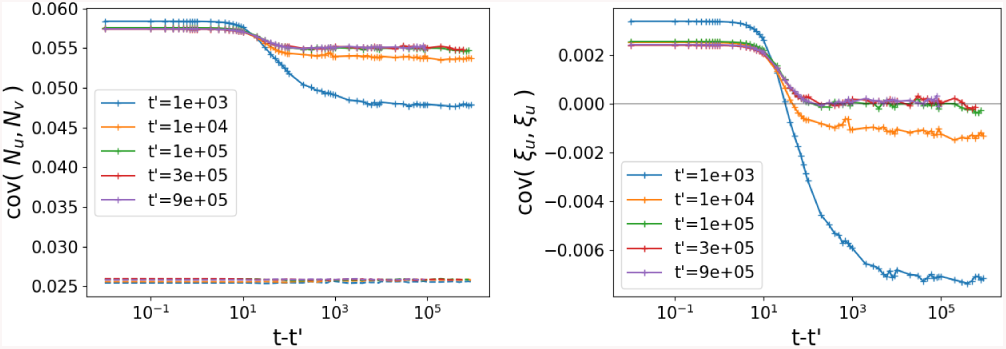
Covariance of the abundances in distinct patches. We use the general notation 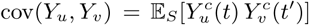 and 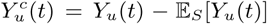. Left: In full lines we show the abundance covariance within a patch *u* = *v*, and across patches *u* ≠ *v* in dotted lines. The correlation in abundances across patches is mainly static: dotted lines are reasonably flat. In other words, the correlation of *ξ*_*u*_ with *ξ*_*v*_ for *u* ≠ *v* is very small. Right: The covariance in *ξ* is shown to reach a TTI state. It only depends on *t* − *t*′ after *t*′ = 10^5^: the colored curves collapse. In this data, 100 distinct simulations were averaged, with parameters (*S, µ, σ* | *M, ρ, d, N*_*c*_) = (400, 10, 2 | 8, 0.95, 10^−10^, 10^−15^).

## Appendix B: DMFT equations

In this section, we present the full DMFT equations, and explain how they can be reduced to the steady-state equations quoted in the main text.

We consider as a starting point equation Eq. (1). For the sake of clarity, we derive DMFT under simplifying assumptions, but the result is much more robust and could be applied to different ecology models as well as real data [17]. DMFT for ecological models has a double valency analogous to the one of mean-field theories in physics: it is at the same time an exact theory for some simple models, and a powerful approximation largely applicable to a broad range of systems. For the sake of clarity, the derivation assumes a fully connected model (all interactions are non-zero), but the results hold for any connectivity *C* as long as *C* ≫ 1, see remark at the end of this Appendix.

The assumptions which make DMFT exact are the following: all constants *N*_*i,u*_(0), *B*_*i,u*_, *D*_*i,uv*_ and *A*_*ij,u*_ are random variables, sampled from known distributions. More precisely:

- in each patch *u* and for all species *i*, the parameters 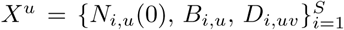 are drawn from a probability distribution ℙ which is a product measure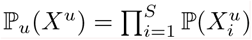;
- the interaction matrix can be decomposed as 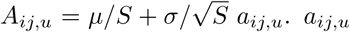 are standard random variables with mean zero, variance one, and correlation:

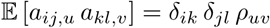

where we used the Kronecker symbol *δ*_*ik*_, and *ρ* _*uv*_ *= ρ* + (1 − *ρ*)*δ*_*uv*_is a uniform correlation *ρ* between patches.

With these conventions, we rewrite Eq. (1) in the following way:

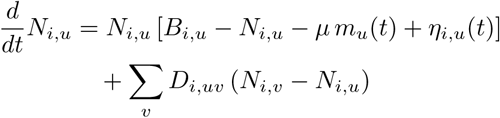

where 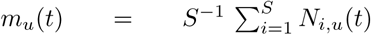 is the mean abundance in patch *u*,and 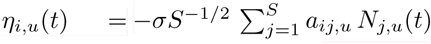 is a species-and-patch-dependent noise.

The DMFT equation can be obtained by following Ref. [14]: in the large-*S* limit, it can be shown that the statistics of this multi-species deterministic process corresponds to the following one-species stochastic process, for each patch.

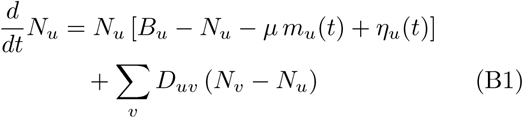

where {*N*_*u*_(0), *B*_*u*_, *D*_*uv*_} are sampled from the distribution ℙ(*X*^*u*^), *m*_*u*_(*t*) is a deterministic function, and *η*_*u*_(*t*) is a zero-mean Gaussian noise. The variability from one species to another becomes in the DMFT setting the randomness contained in {*N*_*u*_(0), *B*_*u*_, *D*_*uv*_} and *η*_*u*_(*t*).

To make this point crystal clear, let us introduce two different averages:

- 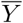 averages over the stochastic process in Eq. (B1): over the stochastic noise *η*_*u*_ and over the distribution ℙ(*X*^*u*^);
- 𝔼_*S*_ (*Y*) denotes the statistical average over the deterministic multi-species system. 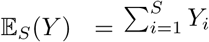,and therefore also includes sampling of 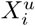.

DMFT represents in terms of a stochastic process the deterministic dynamical system governing the dynamics of the *S* species in the ecosystem. In consequence, averages over the stochastic process coincide with average over species: for a given observable 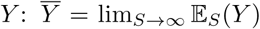. This is analogous to the representation of the environment of an open physical system in terms of thermal noise, as it is done e.g. in the case of the Langevin equation.

The second important aspect of DMFT is *self-consistency*. This is related to the fact that the noise is induced by the dynamics of the species themselves, so its properties can be obtained from dynamical averages:

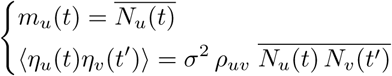

where we used a last average over the stochastic noise only, in order to define its covariance. Henceforth we use the notation 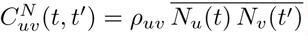.

We now show how DMFT equations simplify for a time-translationally-invariant state of the system, which is in general reached after some transient time. In this state, all one-time observables become constant in time, and two-time observables become functions of the time difference only.

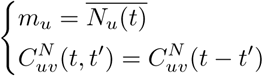

The correlation 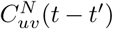 decays at large time differences to a non-zero constant, leading to a static contribution to the noise term. In order to disentangle the static part and the time-fluctuating part of the noise, we perform the decomposition *η*_*u*_(*t*) = *z*_*u*_ + *ξ*_*u*_(*t*) such that *z*_*u*_ and *ξ*_*u*_(*t*) are independent zero-mean Gaussian variables and processes verifying:

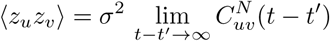

and subsequently 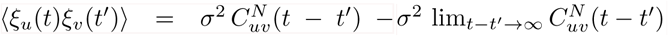 which vanishes for *t* − *t* ′ → ∞. Substituting this decomposition into Eq. (B1), we obtain:

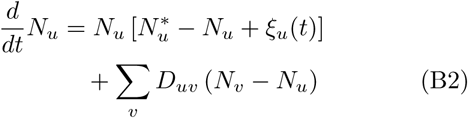

where 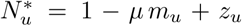 is a Gaussian variable, whose statistics is described in Appendix D. We checked numerically that for small migration *D*, the noise is only correlated between patches through its static part: for *u* ≠ *v, ξ*_*u*_(*t*) *ξ*_*v*_(*t*′) ≪ *z*_*u*_ *z*_*v*_, as presented in Fig. 7. In this case, we can write the self-consistent closure as follows:

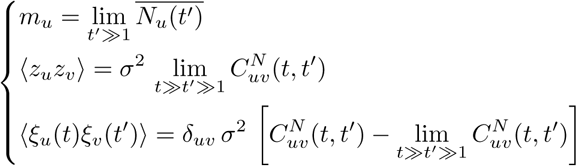

As explained above, DMFT can be implemented as an approximation for a large variety of systems. In this case one has to infer the average *µ*, the standard deviation *σ* of interactions, and the distribution ℙ(*X*^*u*^) from the data (we remind that *X*^*u*^ = {*N*_*u*_(0), *B*_*u*_, *D*_*uv*_}) and use them as an input to define an effective model. The generalization to patch-dependent cumulants *µ*_*u*_ and *σ*_*u*_ is quite straightforward. So is the generalization to patch-dependent correlation *ρ*_*uv*_.

We have derived DMFT for a completely connected set of interactions *A*_*ij*_. A different way to obtain DMFT is considering a finite connectivity network of interactions *A*_*ij*_, e.g. the one produced by a Erdos-Renyi random graph with average connectivity per species *C* or a regular random graph with connectivity *C*. In these cases, for each link *ij* one generates a random variable with average *µ/C* and variance *σ*^2^*/C* and set it to *A*_*ij*_. In the large connectivity limit, *C* → ∞, each species interacts with a very large number of species and one can replace the deterministic interaction with an effective stochastic noise, as done for a completely connected lattice. Although the resulting DMFT equations are the same, the two cases are quite different: in the former a species interact with *C* ≪ *S* species whereas in the latter a species interacts with *C* = *S* species. The equivalence of DMFT for completely connected lattices and finite connectivity ones in the *C*→ ∞ limit has been thoroughly studied in physics of disordered systems in the last twenty years [23].

## Appendix C: Extinction probability of a species

Here the probability of extinction of a species is presented, at the limit *N*_*c*_ ≪ *D* ≪ 1. More specifically, we assume that *N*_*c*_ is small compared to the typical fluctuations of the abundances. In addition, in simulations we see that it is reasonable to assume complete lack of synchrony, namely that the noise *ξ*_*u*_ is uncorrelated between different patches, see Appendix B, Fig. 7. We will therefore assume that in the following calculation. Finally, we assume that for at least one patch, 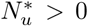, otherwise the species goes quickly extinct.

Within DMFT, the problem thus becomes ones of calculating the extinction probability of a meta-population (single species), under environmental fluctuations, that are uncorrelated between the different patches. We only present the result here; a full account will be given elsewhere.

Let *x*_*u*_ ≡ ln *N*_*u*_. The equations of the DMFT, Eq. (B1), become

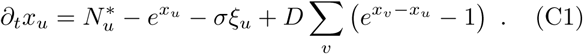

Here *D* = *d/*(*M* − 1). We look for a rare realization of {*ξ*_*u*_} that makes all the *x*_*u*_ reach *x*_*c*_ = ln *N*_*c*_, in the case when the cut-off is low, *x*_*c*_ → −∞. The calculation proceeds within the framework of large-deviation theory [24]. First, one defines the “action”

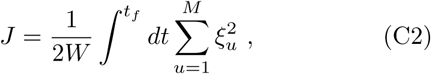

with *ξ*_*u*_ substituted with its value from Eq. (C1), and *W* defined as in the main text. Here we approximated the noise correlations by white noise, which is justified here as the extinction event takes a time which is long compared to the correlation time. We assume that *D* is small.

Then the mean time to the occurrence of such an event scales as 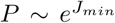 with *J*_*min*_ the action *J* minimized over all population trajectories {*x*_*u*_ (*t*)}*u*=1..*M* that start at *t* → −∞at the typical value of *x*_*u*_, obtained by the zero-noise fixed point of Eq. (C1), and terminate at *t*_*f*_ at *x*_*c*_ = ln *N*_*c*_.

We first describe the result for *M* = 1. In this case there is only one patch, *u* = 1, with 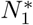. If 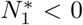 the species is extinct. On the other hand, if 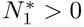, then we obtain the known result [11, 25]

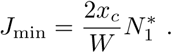

The result for all *M* is a generalization of this result, of the form

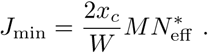

To describe the calculation of 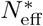, order the patches so that 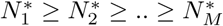. Then there exists 1 ≤ *m* ≤ *M* such that

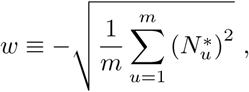

and where *w* satisfies: 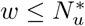 for all *u* ≤ *m*, and 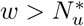 for all *u* > *m*. Such a partition can be shown to always exist. Then

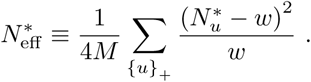

The derivation will be given elsewhere. We illustrate the result by considering two cases. First, in the *M* = 1 example, since 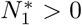, the partition is {*u*}_+_ = {1} and {*u*}_−_ the empty set. Indeed, this gives 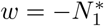, so *w* ≤ 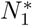. Then 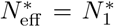 and 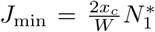, so the result for *M* = 1 is reproduced. Another simple case is when there are *M* patches with identical carrying capacities 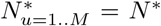. Here {*u*}_+_ = {1, ..*, M*} and *w* = −*N*\*.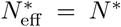, and 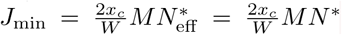. This result is intuitively clear: to go extinct, the species must go extinct in all patches at once, so the probability is 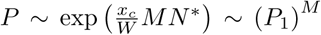, where *P*_1_ is the result for *M*= 1

## Appendix D: Diversity and stability at low migration rates

We use notations from Appendix B. Within the time-translational-invariant state:

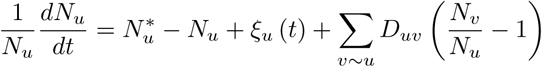

Consider the case of low migration, *D* → 0^+^. We now develop a theory assuming that the amplitude of the en-dogenous fluctuations,

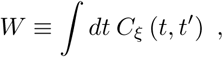

remains finite in the limit *D* → 0^+^. Assume the species survives, i.e. there is at least one patch with 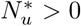. If 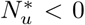 then *N*_*u*_ = *O* (*D*). If 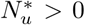 then *N*_*u*_ = *O* (1) and therefore 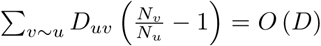. Taking the time average of the above equation

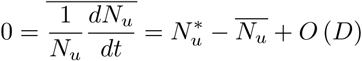

and therefore 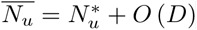.

The previous arguments lead to the conclusion that in the 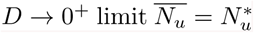 if 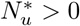 and is equal to zero otherwise. In the following we provide more detail more this argument and its possible limitations. For this last equality to be valid, we need that 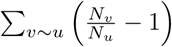 will be finite, so that 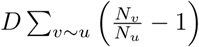 will indeed be small. This might break if *N*_*u*_ can be small while some other *N*_*v*_ remains *O* (1). An estimate for that proceeds by noting that the carrying capacity of patch *u* in the presence of other patches is larger or equal to 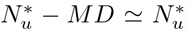, its carrying capacity alone. If patch *u* fluctuates alone, then

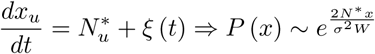

This gives for 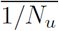

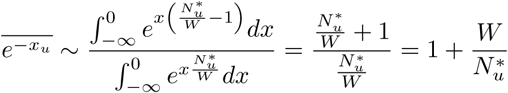

For any given 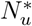 this is finite. It diverges as 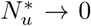. Therefore the migration term is negligible only if 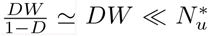.(Note that migration itself would limit *N*_*u*_ going below much below *DN*_*v*_, which would make this term smaller.) The main approximation (or limitation) of our approach is the assumption that *W* remains finite in the small *D* limit. This is shown to hold in simulations presented in Appendix B. It breaks down if the noise develops long-lasting correlations in time. Our approximation will be nevertheless good for large 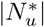 and for weak endogenous fluctuations.

<u>W</u>e now used the relationship discussed above between 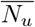 and 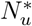 to determine the statistics of 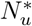. We shall use the term “source” for patches where 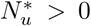, and “sink” otherwise^7^. In order to understand the correlation between the sources in the different communities, we unpack 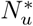 using Appendix B.Taking the time-average is equivalent to averaging over the dynamical noise *ξ*. Therefore, in patch *u* for species 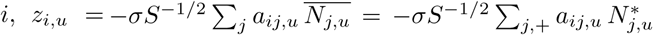. The Σ_*j*,+_ sum means that we only sum over 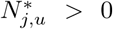 Here, we recall that *a*_*ij,u*_ are standard random variables with mean zero, variance one, and correlation between patches:

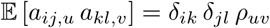

where we used the Kronecker symbol *δ*_*ik*_. Therefore:

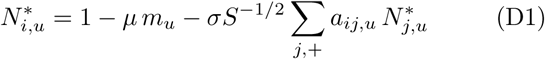

where we recall 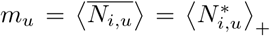. We can now compute the different moments of the multivariate Gaussian random variable 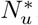, using equation (D1). We obtain the closure:

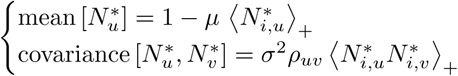

When *u* = *v*, as *ρ*_*uu*_ = 1, we find the expected single community result. In particular, mean 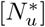 and variance 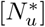 do not depend on the patch *u*.

We numerically solve the closure in a self-consistent way: start with a guess for 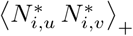, and then (1) Produce many samples of the vector 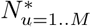 and (2) calculate the next estimate for 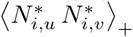, by averaging only over 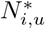 and 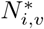 that are both positive. For stability of this numerical scheme, we only replace half the samples at each iteration. We use 10^5^ samples and 1000 iterations. The algorithm is always found to converge to the same solution.

Given covariance 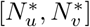, the distribution of 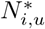 is completely specified: it is a multivariate Gaussian in *u*, has the single-patch statistics of a single community, and a known covariance between patches. The solution can then also give the distribution of the number of sourcing patches.

In addition, we can compute the correlation coefficient *ρ*_*N*_*\** of the 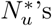. We use here our simple case of a uniform correlation *ρ*_*a*_ between patches *ρ*_*uv*_ = *ρ*_*a*_ + (1 − *ρ*_*a*_)*δ*_*uv*_. We introduce the notation *ρ*_*a*_ instead of ‘*ρ*’ in this section in order to avoid confusion with *ρ*_*N*\*_.

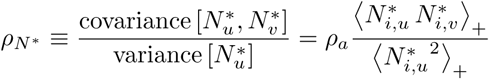

The results are surprising: even when *ρ*_*a*_ *→* 1, the overlap between communities is not perfect (*ρ*_*N*_*\* <* 1), so the total diversity is larger than the one in each patch. Th<u>i</u>s happens *exactly at* the transition to chaos at 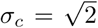, see Fig. 8.

**Figure 8.**
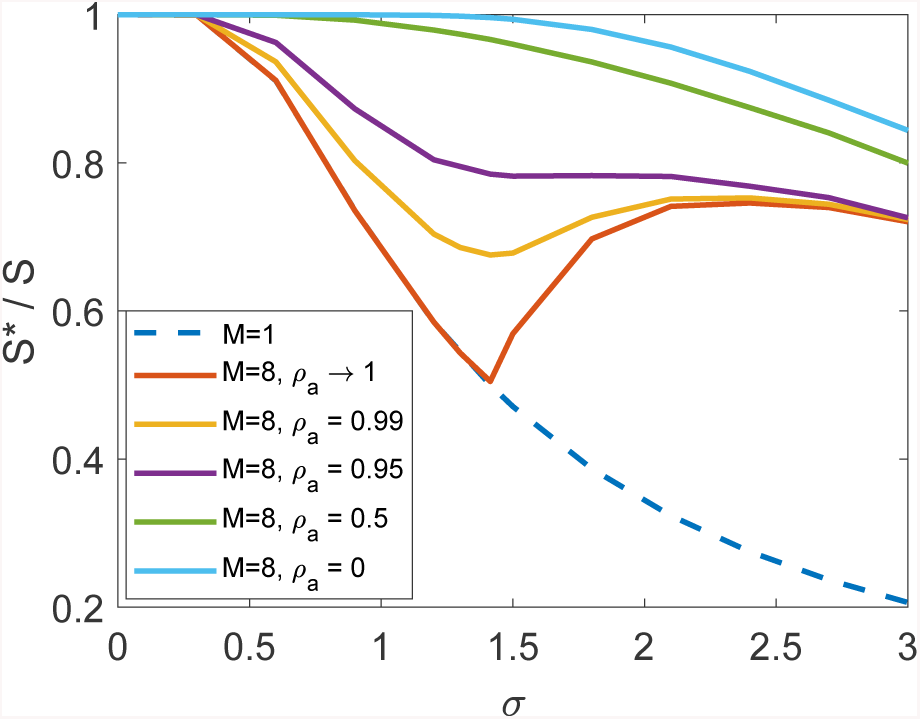
Theoretical predictions for the diversity as a function of *σ* for *M* = 1, 8 patches, *ρa* = 0, 0.5, 0.95 and *ρa →* 1

On Fig. 10, we compare the theory predictions to simulations. In terms of diversity, the theory appears to give an upper bound to the simulations. The difference becomes larger at higher values of *σ*, and for *ρ*_*a*_ closer to one. To look further into this difference, it is useful to study diversity as a function of the value of 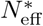 of each species. As shown in Fig. 6 in the main text, most of the difference in diversity is due to low values of 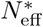, which are precisely the species that are more likely to go extinct, with good agreement with theory at higher values of 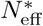. This is demonstrated in Fig. 9, which shows that the theoretical prediction for the number of species with 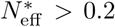 is closer to simulation results than the predictions for total diversity. At the moment we do not know if remaining differences are because the theoretical argument is only approximate, or whether in principle, with exceedingly low values of *N*_*c*_ and *D*, it could be approached by simulations for any *σ*.

**Figure 9.**
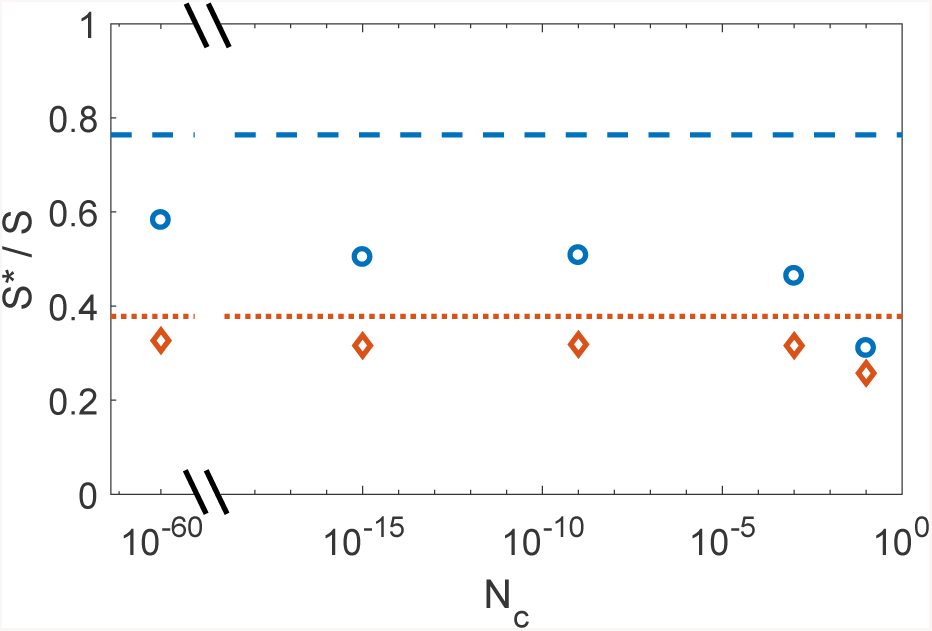
The fraction of persistent species *S*^*\**^*/S* (circles) is compared to theoretical bound (blue dashed line), for different values of *N*_*c*_. Also shown is the fraction of species above 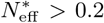, compared to the theoretical bound for that (red dotted line), showing better agreement than for the full diversity. Simulations use the same parameters as in Fig. 6, but with a range of values for *N*_*c*_ (Fig. 6 corresponds to the points at *N*_*c*_ = 10^−15^).

**Figure 10.**
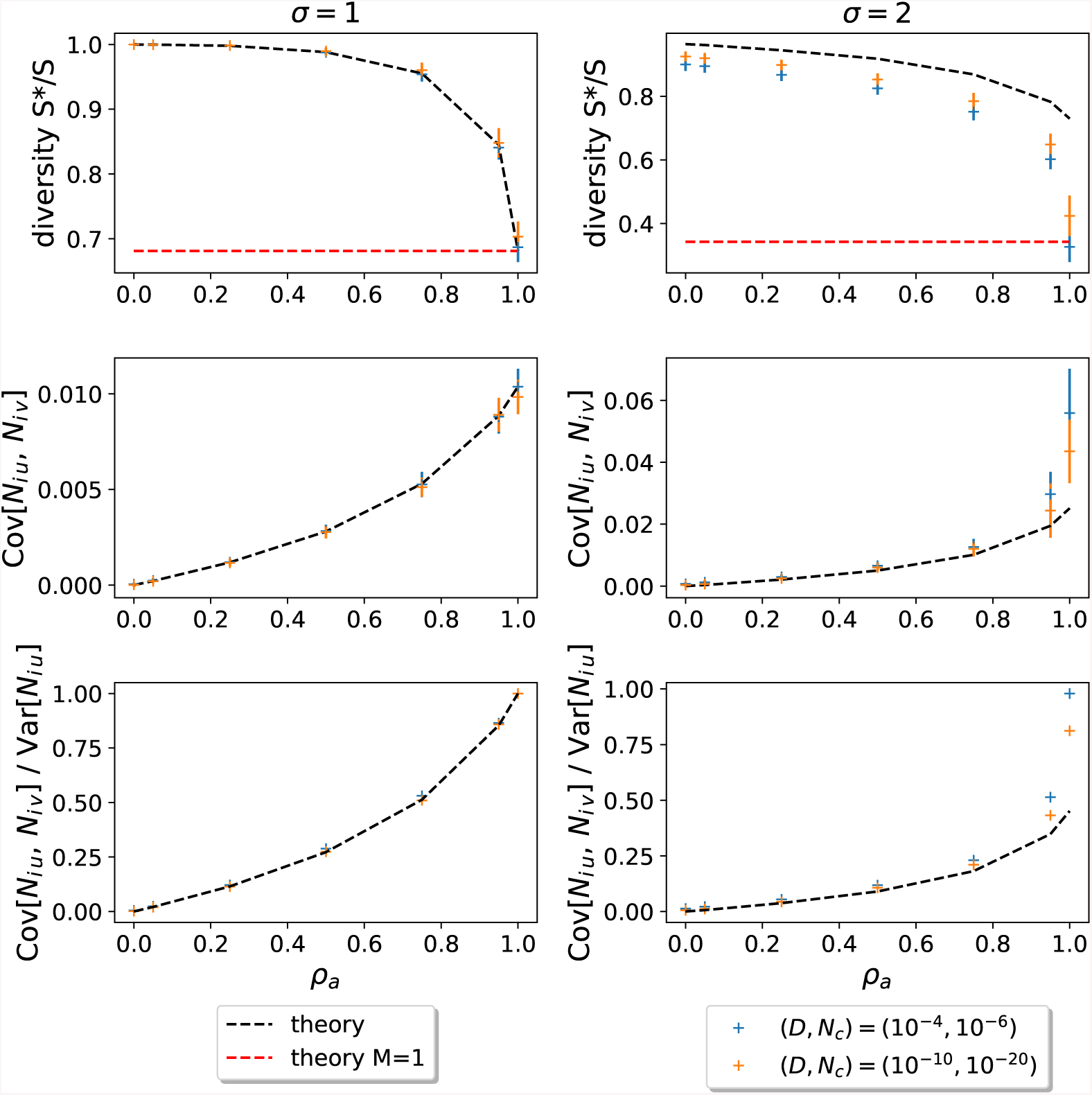
Numerical checks of the theoretical predictions. From top to bottom, we consider three different observables: the diversity, the covariance in the abundances across distinct patches, and this covariance rescaled by the one patch variance. By varying *σ*, we can control the state of the system: on the left (*σ* = 1), we show the results for fixed points; on the right 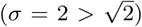, we show the results for persistent dynamical fluctuations. In dotted lines, we plot the theory predictions, as functions of the correlation between patches’ interactions *ρ*_*a*_. We compare them to simulations with parameters (*S, M, μ*) = (400, 8, 10), and obtained by simulations run until final time *t*_*f*_ = 10^4^. We eventually vary the couple (*D, N*_*c*_). We use 50 distinct samples of the simulations for each combination of parameters, in order to get error bars and relevant statistics. The cut-off is implemented via patch-wise extinctions when the abundance goes below the threshold in each particular patch, in which case migration out of the patch is turned off while still allowing inward migrations. On the left side, we can see that the theory is exact in the fixed point regime. In this regime, as *ρ*_*a*_ →1, the predictions are equivalent to the one patch *M* = 1 theory, as all patches are the same. In the persistent fluctuation state, the theory is only a good approximation. More precisely, the predictions become more accurate as *D* and *N*_*c*_ go to zero, as expected. In addition, the agreement gets worse when *ρ*_*a*_ →1, because synchronization can occur. In the top right figure, we show that the prediction for diversity is an upper bound. In the bottom right figure, we see that indeed the prediction for *ρ*_*n*_ is still far from 1 when *ρ*_*a*_ → 1, for the values of *D, N*_*c*_ used in the simulations.

To find the boundary of parameter space where fixed points loose their stability and the system becomes chaotic, we look at the linear stability of persistent species.When *D* is small, the species that are not sourced in each patch do not affect the stability, and so the question simplifies to single patch stability, which when corr [*A*_*ij*_, *A*_*ji*_] = 0, results in 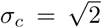 and with 1*/*2 of the species being sourced in each patch [26].

## Appendix E: Single patch (*M* = 1)

Here we show that in principle a single patch can reach and maintain a dynamically fluctuating state. However, this requires prohibitively large *S*, not attainable in practice. In Fig. 11 and Fig. 12 we show results of a numerical solution [14] to the DMFT equations detailed in Appendix B. At extremely low values of *N*_*c*_ the system appears to reach a final diversity above the May bound and, hence, to be chaotic. DMFT however describes the behavior in the *S* ≫ 1 limit. When full simulations of the model in Eq. (1) are carried out at finite *S*, they diversity falls somewhat below the DMFT final diversity, leading to a fixed point, rather than a chaotic state, see Fig. 12. This finite-size correction to the DMFT result are important since they show that maintaining a dynamically fluctuating state for realistic values of *S* is not possible for *M* = 1.

**Figure 11.**
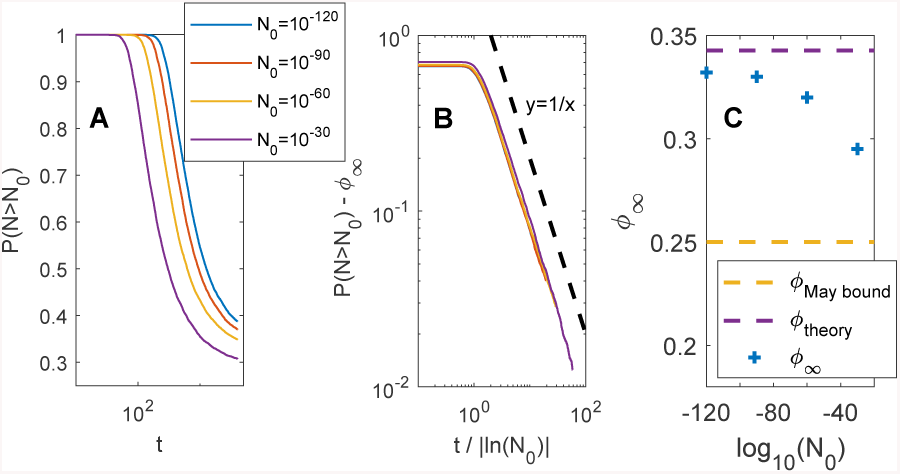
DMFT numerics for a single patch, *M* = 1, showing that chaos is in principle possible here, although for unrealistic values of model parameters. (A) The fraction of species above different values of *N*_0_, *P* (*N* > *N*_0_) is plotted as a function of time, for different values of *N*_0_. (B) The curves for different *N*_0_ collapse when *P* (*N* > *N*_0_) −*ϕ*_∞_ (*N*_0_) ∼ |ln *N*_0_| */t*. Here *ϕ*_∞_ (*N*_0_) is a fitted parameter, the extrapolated value of *P* (*N* > *N*_0_) at long times. (C) The values of *ϕ*_∞_ (*N*_0_) are well above the linear stability bound (“May bound”), and at (very) low *N*_0_ come quite close to the theoretical maximal value for *ϕ*_∞_ (*N*_0_), predicted in Appendix D.Here *σ* = 2, *μ* = 10, *N*_c_ = 10^−120^.

**Figure 12.**
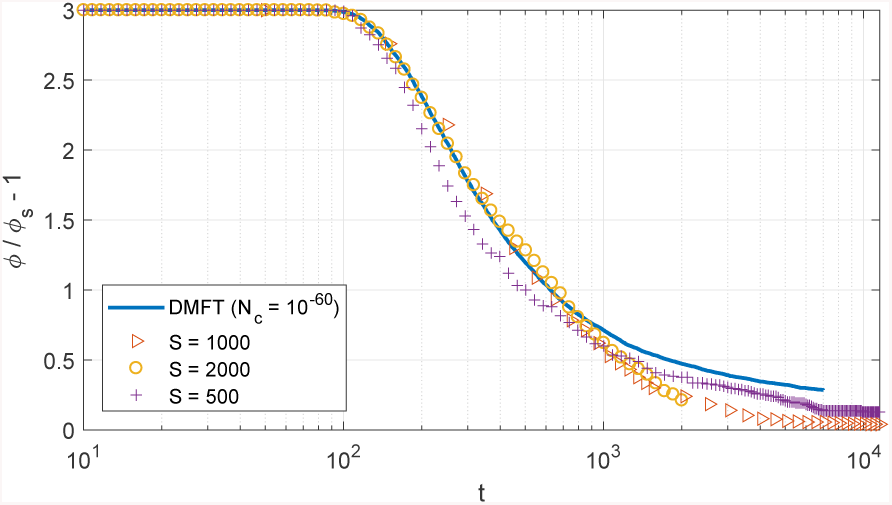
The DMFT solution and the simulations only agree up to times *t* ∼10^3^, after which the diversity in the simulations reduces more rapidly and reaches a fixed point. This means that the convergence to the DMFT solution is slow with *S*.

## Appendix F: Correlations of interactions in a pair of species

In the main text we assumed that *A*_*ij,u*_ is sampled independently from *A*_*ji,u*_. Here we show that the long-lived endogenous fluctuations can be found even if this assumption is relaxed. For this purpose, we consider a symmetric network of non-zero *A*_*ij,u*_, namely *A*_*ij,u ≠*_ 0 if and only if *A*_*ji,u*_. We define *γ* the correlation of the non-zero elements 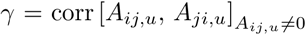. Fig. 13 shows two simulations, one with *γ >* 0 and the other with *γ <* 0. In both cases the system relaxes to a long-lived state with fluctuating abundances, without further loss of diversity up to time 2 · 10^5^. They are intended solely to demonstrate that conditions with *γ* ≠ 0 exist, rather than a systematic exploration of such cases.

**Figure 13.**
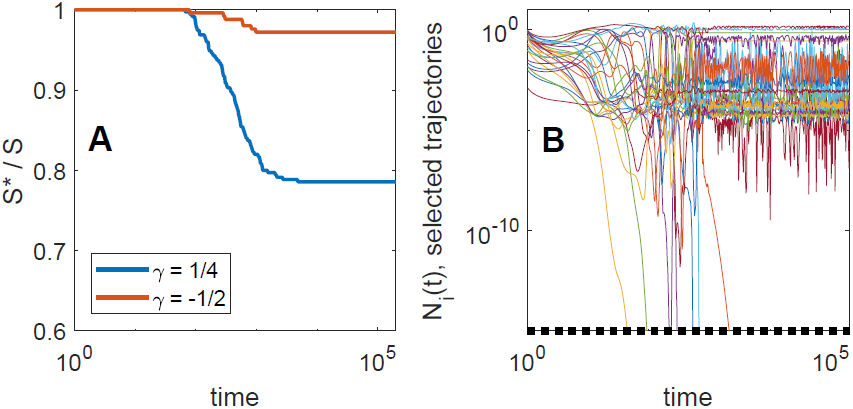
(A) The diversity *S*^*^ (*t*) */S* for two runs with *γ* ≡ corr [*A*_*ij,u*_, *A*_*ji,u*_] ≠ 0. (B) Selected trajectories of *N*_*i,u*_ (*t*) for the run with *γ* = 1/4.

The parameters for the simulations (using the notation of Appendix A) are the following: Run with positive *γ*: *γ* = 1*/*4, *S* = 350, mean (*A*_*ij,u*_) = 0.075, std (*A*_*ij,u*_) = 0.175, *c* = 0.357, *M* = 8, *d* = 10^−3^, *ρ* = 0, *N*_*c*_ = 10^−15^.

Run with negative *γ*:*γ*= −1*/*2, *S*= 250, mean (*A*_*ij,u*_) = 0.075, std (*A*_*ij,u*_) = 0.358, *c* = 0.5, *M* = 8, *d* = 10^−3^, *ρ* = 0, *N*_*c*_ = 10^−15^.

## Footnotes

1 We are interested in the regime where recolonization by migration between patches is fast compared to the rate of extinction events. In this regime, we expect (and checked in a few cases) that other implementations of the cut-off *N*_*c*_ will lead to the same qualitative phenomena. For instance, we implemented patch-wise extinctions when the abundance goes below the threshold in one particular patch, while still allowing migrations in.

2 In the main text we focus on the asymmetric case in which *A*_*ij,u*_ and *A*_*ji,u*_ are uncorrelated. We show in Appendix F that our results also hold when correlations are present.

3 Each interaction coefficient is non-zero with probability *c* = 1*/*8, and the non-zero interactions are Gaussian with mean (*A*_*ij,u*_) = 0.3, std (*A*_*ij,u*_) = 0.45.

4 We will see in the next section that, when endogenous fluctuations are present, this state is actually metastable, i.e. it is almost stationary on large but finite time-scales.

5 It verifies the self consistent equation: 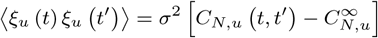 Where 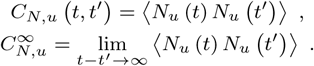 The average ⟨..⟩ over the stochastic process corresponds to the average over species in the original Lotka-Volterra equations.

6 The expresson of the time scale is analogous to the Arrhenius law for activated processes in physics and chemistry: in this case, the counterpart of the energy barrier is −[*N*\* ln *N*_*c*_] and fluctuation amplitude *W* plays the role of the temperature.

7 The term “source” is used here so as to include patches (some-times referred to as pseudo-sinks) where a species might still receive migration from patches with even larger 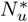. But the contribution of this migration is small and not required for its persistence.

